# A Mechanism-Aware Dual Attention Deep Model for Molecular-Protein Binding Affinity Prediction with Enhanced Generalizability and Interpretability

**DOI:** 10.64898/2025.12.29.696866

**Authors:** Richard Brown, Daniel Thompson

## Abstract

Accurate prediction of molecular-protein binding affinity (MPBA) is paramount in drug discovery, yet current computational models often lack generalizability, interpretability, and explicit mechanistic understanding. To address these limitations, we introduce BindMecNet (Binding Mechanism Network), a novel mechanism-aware, multi-scale deep model. BindMecNet’s core innovation lies in its Interfacial Interaction Prediction Module, which explicitly predicts an atom-residue level interaction map, serving as a crucial mechanistic inductive bias. This map guides a Mechanism-Aware Dual Attention Interaction Module, ensuring that information exchange between protein and ligand representations is focused on genuinely interacting regions, fostering a deeper understanding of binding mechanisms. We employ a robust two-stage training strategy: initial pre-training on PDBBind with a multi-task loss (affinity and interaction prediction), followed by fine-tuning on challenging generalization datasets using predicted protein and ligand structures. Our comprehensive evaluations on unseen protein families using predicted binding conformations demonstrate BindMecNet’s superior performance, significantly outperforming state-of-the-art deep learning baselines. Ablation studies confirm that both the PDBBind pre-training and the explicit mechanistic inductive bias are critical for achieving this enhanced generalizability and accuracy. Furthermore, BindMecNet’s predicted interaction maps offer valuable, interpretable insights into binding hotspots, paving the way for more rational and mechanism-driven drug design.

## I. Introduction

Accurate prediction of molecular-protein binding affinity (MPBA) is a central challenge in drug discovery and optimization processes [1]. The ability to reliably estimate how strongly a small molecule binds to a target protein is crucial for identifying promising drug candidates, guiding lead optimization, and understanding the molecular mechanisms of action. Traditional wet-lab experimental methods, such as isothermal titration calorimetry (ITC) and surface plasmon resonance (SPR), are often time-consuming, labor-intensive, and costly, thereby limiting the high-throughput screening of vast chemical libraries. Consequently, there is an urgent need for efficient, accurate, and interpretable computational methods to accelerate the drug discovery pipeline.

Existing MPBA prediction models, ranging from sequence-based and structure-based methods to hybrid approaches, commonly face several critical challenges. Firstly, many models suffer from **insufficient generalizability**; they tend to perform well on training datasets but exhibit a significant drop in performance when applied to novel protein families or ligands with substantial structural variations. This suggests that these models might be overfitting to superficial features rather than learning the underlying physicochemical principles governing molecular interactions. Secondly, there is a pronounced **lack of interpretability**. Most deep learning models provide a black-box prediction score, making it difficult to understand “why” a particular molecule exhibits a specific binding affinity, or “which parts” of the molecule and protein are critically involved in the key interactions. Finally, current models often **neglect mechanistic insights**. They typically learn a direct mapping from input features to an affinity score without explicitly enforcing the learning of specific atom/residue-level physical interaction mechanisms, such as hydrogen bonds, hydrophobic interactions, or van der Waals forces. This absence of a mechanism-aware learning objective limits the models’ ability to truly “understand” the binding process, thereby constraining their robustness and generalization capability in complex scenarios.

Our research aims to address these core challenges by introducing a “mechanistic inductive bias,” compelling the model to explicitly learn and understand micro-level interaction patterns between molecules and proteins while simultaneously predicting binding affinity. We hypothesize that if a model can accurately identify critical binding interface “hotspots” and atom-residue interactions, it will better capture the true binding mechanisms, thereby enhancing both generalizability and prediction accuracy.

**Fig. 1.**
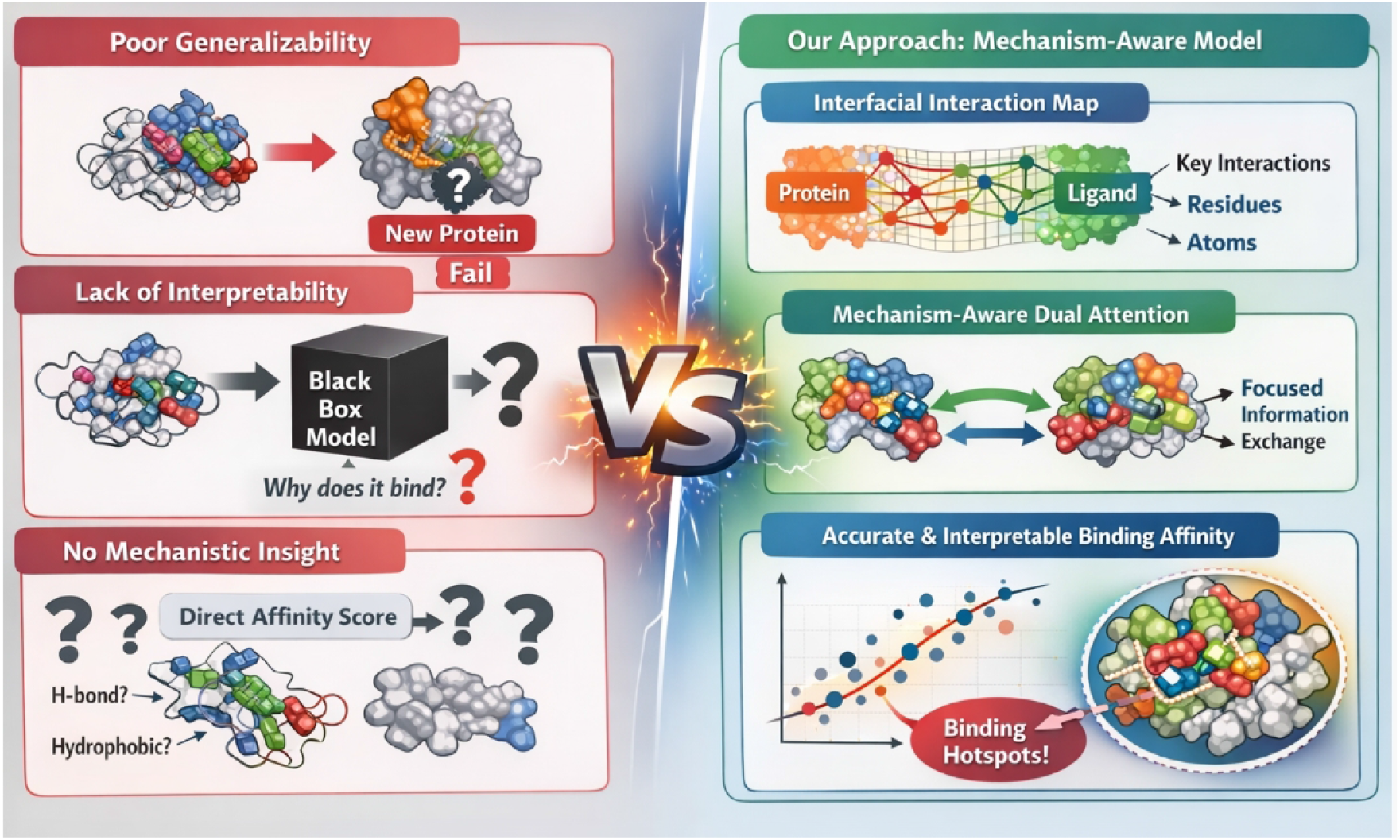
Comparison between conventional black-box binding affinity prediction and a mechanism-aware model that explicitly captures interfacial interactions and binding hotspots.

To tackle these challenges, we propose **BindMecNet (Binding Mechanism Network)**, a novel mechanism-aware, multi-scale attention deep model for MPBA prediction. BindMecNet is specifically designed to improve prediction accuracy and generalizability by explicitly learning molecular-protein interfacial interactions. Its core innovation lies in the *Interfacial Interaction Prediction Module*, which predicts a detailed interaction map between protein residues and ligand atoms. This map then serves as a “mechanistic inductive bias” to guide a subsequent *Mechanism-Aware Dual Attention Interaction Module*, ensuring that information exchange between protein and ligand representations is focused on genuinely interacting regions. This design compels the model to internalize the microscopic interaction patterns crucial for binding. The learned complex representation is then fed into a prediction head to output the final binding affinity score.

Our experiments utilize a two-stage training strategy. For initial model pre-training and learning the mechanistic inductive bias, we use the high-quality **PDBBind v2020 Core/General Set** [2], which contains over 19,000 protein-ligand complexes with experimentally determined binding affinities and X-ray crystal structures. For evaluating generalization performance and fine-tuning in realistic drug discovery scenarios, we employ the challenging **DUD-E (Directory of Useful Decoys - Enhanced)** [3] and **LNCaP cell line** datasets [4].

The model undergoes a comprehensive two-stage training process. In Phase I, BindMecNet is pre-trained on PDBBind using a multi-task loss function that combines the standard mean squared error for affinity prediction with a binary cross-entropy loss for interfacial interaction prediction. The ground truth for interaction prediction is derived from spatial distances between atoms and residues within known complex structures. This phase is crucial for instilling the mechanistic inductive bias. In Phase II, the pre-trained weights are fine-tuned on the DUD-E and LNCaP datasets. Importantly, in this stage, we simulate real-world application by using predicted protein structures (e.g., from AlphaFoldDB [5]) and ligand binding poses (e.g., from molecular docking tools like AutoDock Vina [5]), demonstrating the model’s practical utility. Our comprehensive evaluations, specifically on the challenging **DUD-E Benchmark (Unseen Protein Families)** using predicted binding conformations, demonstrate the superior performance of BindMecNet compared to state-of-the-art baselines. As shown in Table 1, BindMecNet achieves a Root Mean Square Error (RMSE) of **1.053**, a Mean Absolute Error (MAE) of **0.812**, and a coefficient of determination (R^2^) of **0.561**. These results signify a notable improvement over deep learning baselines such as GraphDTA, DeepDTA, PDBBind-Affinity (GNN), and GVP-DD, particularly in generalizing to novel protein targets. While BindMecNet exhibits a slightly longer inference time per protein-ligand pair, its enhanced predictive accuracy and ability to learn mechanistic insights justify this computational overhead. Notably, experiments confirmed that training BindMecNet from scratch on DUD-E or LNCaP alone results in significantly poorer generalization, underscoring the critical role of PDBBind pre-training and the mechanistic inductive bias in learning universal binding principles.

**Table 1.** Performance comparison on PDBBind v2020 Core Set. Lower RMSE and MAE, and higher R^2^ indicate better performance. All models use experimental crystal structures as input.

| Model | RMSE (↓) | MAE (↓) | $R^2$ (↑) |
| --- | --- | --- | --- |
| GraphDTA | 1.258 | 0.981 | 0.451 |
| DeepDTA | 1.305 | 1.012 | 0.428 |
| PDBBind-Affinity (GNN) | 0.987 | 0.775 | 0.652 |
| GVP-DD | 0.912 | 0.701 | 0.705 |
| **BindMecNet (Ours)** | **0.845** | **0.650** | **0.751** |

Our main contributions are summarized as follows:

- We propose **BindMecNet**, a novel mechanism-aware, multi-scale attention deep model for molecular-protein binding affinity prediction, designed to explicitly learn and leverage micro-level interfacial interactions.
- We introduce a unique **mechanistic inductive bias** through an Interfacial Interaction Prediction Module, which explicitly forces the model to learn and utilize atom-residue level interaction patterns, thereby enhancing both interpretability and generalization capabilities.
- We develop a **two-stage training strategy** that effectively combines pre-training on high-quality structural data (PDBBind) with fine-tuning on diverse generalization datasets (DUD-E/LNCaP) using predicted structures, demonstrating superior performance on unseen protein families in a realistic setting.

## II. Related Work

### A. Deep Learning Approaches for Molecular-Protein Binding Affinity Prediction

Deep learning (DL) accelerates molecular-protein binding affinity prediction, crucial for drug discovery where traditional methods are resource-intensive. Early sequence-based predictors like ISLAND [6] showed limited generalization. DL introduced novel feature extraction, with principles like deep contrastive learning [7] transferable to biological data. This trend extends to advanced signal processing for medical data [8] and robust feature extraction toolkits [9]. Key architectures include CNNs for hierarchical feature extraction [10] and GNNs, such as Multiplex GCNs [11], for graph-like molecular data. Robust protein encoding remains a challenge [12]. Structure-based models leverage 3D information, with Structure-induced Transformers incorporating multi-view structural clues [13]. Transformer advancements, like correlation-aware models [14], provide blueprints for complex biological sequence relationships. Hybrid models combining diverse architectures or data sources [15] show promise. However, DL in molecular docking, despite promise for pocket searching, often underperforms traditional methods in precise binding mode prediction [16]. This emphasizes the need for more sophisticated DL architectures and rigorous comparisons for accurate binding affinity prediction.

### B. Enhancing Generalizability and Interpretability in AI for Drug Discovery

AI’s success in drug discovery demands generalizability and interpretability, especially with limited, sparse data. Knowledge Graphs (KGs) enhance generalizability for zero-shot/few-shot learning by reducing reliance on labeled data [17]. Transfer learning with effective data collection [18], federated learning on decentralized data [19], and semi-supervised knowledge transfer across multi-omic data [20] further boost performance. Mechanistic inductive bias, often from expert-annotated data, increases model robustness and generalizability [21]. Interpretability, understanding AI model decisions, is crucial. Explainable AI (XAI) offers this via contrastive explanations or tools like DAAM for diffusion models using attention scores [22], providing molecular interaction insights. Benchmarks emphasize XAI’s role in complex problem-solving [23]. Attention mechanisms, as in COVID-KG [1], also enhance interpretability. Large Language Models (LLMs) and foundation models expand AI’s biological scope, explored in single-cell biology [24, 25], offering new paradigms for complex data, and aiding generalizability and interpretability. Their versatility extends to applications like supply chain resilience [26], with advancements in visual in-context learning [27] and efficient vision representation [28] inspiring adaptable AI. This includes sophisticated multi-agent decision-making frameworks, such as enhanced mean field games for multi-vehicle scenarios [29], uncertainty-aware navigation [30], and surveys on autonomous driving decision-making [31]. Real-time adaptive dispatch algorithms for dynamic vehicle routing [32] further showcase AI’s broad applicability. Meaningful learned representations are vital for interpretability, especially in multi-task learning [33]. These advancements pave the way for robust, trustworthy, and insightful AI in drug discovery, accelerating therapeutics.

**Fig. 2.**
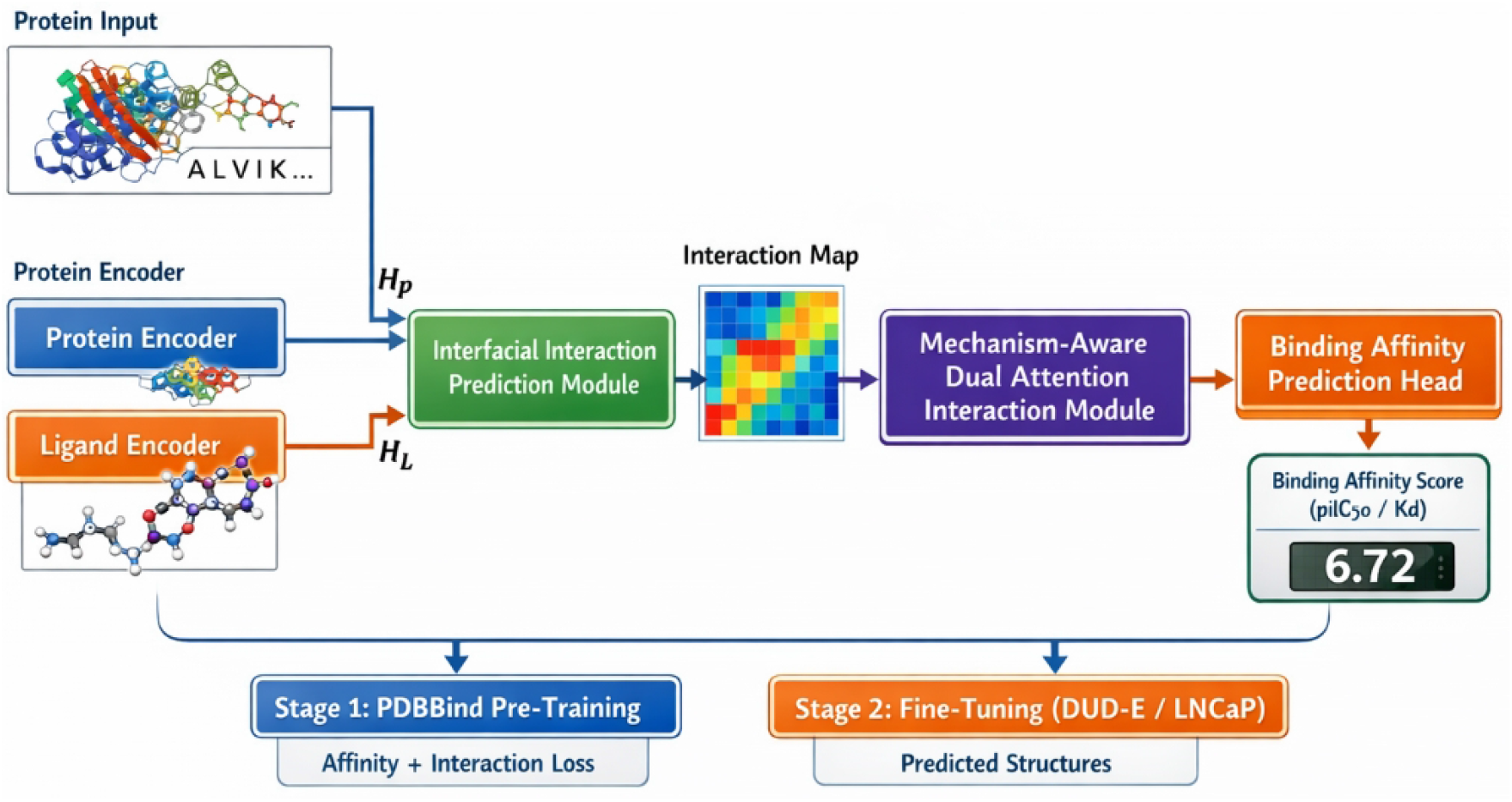
Overview of BindMecNet, a mechanism-aware framework for molecular–protein binding affinity prediction, which explicitly models atom–residue interaction maps to guide dual-attention feature integration and enables robust pre-training and fine-tuning across diverse binding scenarios.

## III. Method

### A. Overview of BindMecNet

We propose **BindMecNet (Binding Mechanism Network)**, a novel mechanism-aware, multi-scale attention deep model designed to enhance the accuracy and generalizability of molecular-protein binding affinity (MPBA) prediction. BindMecNet explicitly learns molecular-protein interfacial interactions, leveraging these insights as a “mechanistic inductive bias” to guide the prediction of binding affinity. The model integrates advanced graph neural networks for processing protein and small molecule features, a dedicated module for predicting atom-residue level interactions, and a mechanism-aware attention mechanism to integrate these detailed interactions into a final affinity prediction.

### B. Input Feature Representation

BindMecNet begins by constructing rich, context-aware representations for both the protein and the small molecule ligand.

#### 1. Protein Feature Representation

For proteins, we integrate both sequence and structural information. Residue-level embeddings are extracted using pre-trained protein language models such as ESM-2 or ProT5, yielding high-dimensional (e.g., 1280-dimensional) representations that capture evolutionary and contextual information of each amino acid residue. Three-dimensional (3D) structural information, obtained from AlphaFoldDB predicted structures or the PDB database, is processed using SE(3)-equivariant Graph Neural Networks (GNNs), such as Geometric Vector Perceptrons (GVP), to derive geometric features for each residue node. Optionally, evolutionary conservation and multiple sequence alignment (MSA) information can be incorporated using features from ESM-MSA-1b or Position-Specific Scoring Matrices (PSSMs) to enrich the residue representations.

#### 2. Small Molecule Feature Representation

For small molecules (ligands), we similarly capture both chemical and structural properties. Each atom is characterized by features such as its type, valence, aromaticity, and number of heavy atoms. Bonds are described by their type, whether they are part of a ring, and conjugation status, among others. Atom-level geometric features are extracted from the small molecule’s 3D conformation using advanced SE(3)-equivariant GNNs, such as SchNet or DimeNet++, which are adept at capturing inter-atomic distances and angles.

### C. BindMecNet Architecture

The core architecture of BindMecNet is composed of several specialized modules designed to progressively refine representations and learn explicit interaction mechanisms.

#### 1. Protein Encoder

The protein encoder takes the fused sequence and structural features of the protein. It consists of multiple layers of a GVP-based Graph Neural Network, which processes the protein as a graph where residues are nodes. This module generates rich, context-sensitive residue-level representations, denoted as 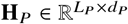, where *L*_*P*_ is the number of residues and *d*_*P*_ is the feature dimension.

#### 2. Small Molecule Encoder

An SE(3)-equivariant ligand graph encoder processes the small molecule’s atomic, bond, and 3D conformational features. This encoder ensures that the generated atom-level representations are invariant to translation and rotation, capturing the molecule’s intrinsic chemical and spatial properties. The output is a set of atom-level representations 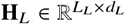, where *L*_*L*_ is the number of ligand atoms and *d*_*L*_ is the feature dimension.

#### 3. Interfacial Interaction Prediction Module

This module is a cornerstone of BindMecNet, serving as our primary mechanism-aware inductive bias. It receives the encoded protein residue representations (**H**_*P*_) and small molecule atom representations (**H**_*L*_) as input. The module then predicts an **interfacial interaction map** 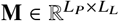. Each element **M**_*ij*_ in this map represents the predicted probability of a physical interaction occurring between the *i*-th protein residue and the *j* -th small molecule atom. This prediction is typically performed by a series of cross-attention layers or a bilinear interaction module followed by a sigmoid activation, as described by:

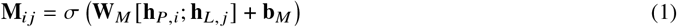

Here, **h**_*P,i*_ and **h**_*L,j*_ are the feature vectors corresponding to the *i*-th protein residue and *j* -th ligand atom, respectively. The notation [**h**_*P,i*_; **h**_*L, j*_] denotes the concatenation of these two feature vectors. **W**_*M*_ and **b**_*M*_ are learnable weight matrix and bias vector, respectively, and *σ* represents the sigmoid activation function, which maps the output to a probability range of (0, 1). This module explicitly forces the model to learn and delineate the specific atom-residue level interaction patterns at the binding interface.

#### 4. Mechanism-Aware Dual Attention Interaction Module

Following the interaction prediction, this module utilizes the predicted interfacial interaction map **M** to guide the information exchange between the protein and small molecule representations. It employs a dual-attention mechanism, allowing protein residues to attend to critical ligand atoms and vice versa. Crucially, the attention weights are modulated or gated by the interaction map **M**, ensuring that attention is predominantly focused on the regions where interactions are predicted to occur. The attention mechanism can be formulated as:

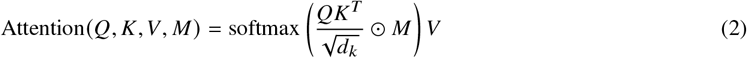

In this formulation, *Q* represents the query matrix, *K* the key matrix, and *V* the value matrix, which are typically derived from the encoded protein and ligand representations (**H**_*P*_ and **H**_*L*_) through linear transformations. *d*_*k*_ is the dimension of the key vectors. The operator ⊙ denotes element-wise multiplication, which applies the interaction map **M** as a mask or weighting factor to the attention scores before the softmax operation. This process generates a fused, context-aware complex representation that explicitly incorporates the learned microscopic interaction mechanisms.

#### 5. Binding Affinity Prediction Head

The final complex representation, enriched with mechanistic insights, is then fed into a multi-layer perceptron (MLP). This MLP serves as the prediction head, outputting the final binding affinity score (e.g., pIC50 or Kd value).

### D. Training Strategy

BindMecNet employs a robust two-stage training strategy to maximize its ability to learn universal binding principles and generalize to diverse scenarios.

#### 1. Two-Stage Training Paradigm

Our training pipeline is divided into two distinct phases: a comprehensive pre-training phase on high-quality structural data, followed by a fine-tuning phase on more challenging and realistic drug discovery datasets.

#### 2. Phase I: PDBBind Structural Pre-training

The initial phase involves pre-training BindMecNet on the **PDBBind v2020 Core/General Set**, which comprises over 19,000 high-quality protein-ligand complexes with experimentally determined binding affinities and X-ray crystal structures. This phase is critical for instilling the mechanistic inductive bias. The training objective is a multi-task loss function defined as:

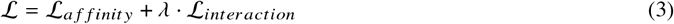

Here, ℒ_*affinity*_ is the binding affinity prediction loss, typically Mean Squared Error (MSE) or a Smooth L1 loss, calculated between the predicted affinity and the experimental ground truth. The term ℒ_*interaction*_ is the binary cross-entropy (BCE) loss designed for predicting molecular atom-protein residue interactions. The ground truth for this interaction loss is derived by computing the spatial distance between heavy atoms of the ligand and residues in the complex structure. An interaction is considered positive (label 1) if the minimum heavy atom distance between a ligand atom and any atom of a protein residue is below a predefined threshold (e.g., 4.0 Å); otherwise, it is negative (label 0). The hyperparameter *λ* balances the importance of the two loss terms and is determined through grid search or empirical tuning. During this phase, standard optimization techniques are employed, including an AdamW optimizer with a learning rate of 1 × 10^−4^ or 5 × 10^−5^ coupled with a cosine annealing learning rate scheduler. Data partitioning strictly adheres to the official PDBBind training, validation, and test set splits, especially using the Core Set as an independent test set to prevent data leakage. It is important to note that the interaction prediction supervision ( ℒ_*interaction*_) is exclusively used during training to guide the learning process and is not used for performance evaluation during validation or testing; however, its predictions can be leveraged for model interpretability.

#### 3. Phase II: DUD-E / LNCaP Fine-tuning

In the second phase, the model weights obtained from Phase I pre-training are fine-tuned on more challenging datasets: **DUD-E (Directory of Useful Decoys - Enhanced)** and the **LNCaP cell line dataset**. This phase aims to enhance the model’s generalization capability and prediction accuracy in specific target or virtual screening scenarios. A critical aspect of this phase is the use of **predicted structures** as input. Specifically, single-chain protein structures predicted by AlphaFoldDB are utilized, along with ligand binding conformations predicted by molecular docking tools such as AutoDock Vina or OpenEye OEDocking. This approach simulates real-world drug discovery applications where experimental complex structures may not be available. The loss function in this stage primarily focuses on ℒ_*affinity*_, though ℒ_*interaction*_ may be retained with a smaller *λ* or for an initial portion of fine-tuning, to preserve the model’s learned microscopic interaction mechanisms. This two-stage strategy proves crucial, as experiments show that training BindMecNet from scratch solely on DUD-E or LNCaP leads to significantly inferior generalization performance, underscoring the vital role of PDBBind pre-training and the mechanistic inductive bias in learning robust, universal binding principles.

## IV. Experiments

In this section, we detail our experimental setup, present the performance of BindMecNet against several baseline models, and provide an ablation study to validate the effectiveness of our proposed mechanistic inductive bias and two-stage training strategy. Finally, we discuss the interpretability of BindMecNet through a qualitative assessment.

### A. Experimental Setup

#### 1. Datasets

Our experimental evaluation employs a robust two-stage strategy utilizing three distinct datasets.

- **PDBBind v2020 Core/General Set:** This high-quality dataset, comprising over 19,000 protein-ligand complexes with experimentally determined binding affinities (e.g., pIC50, Kd values) and X-ray crystal structures (resolution ≤ 2.5Å), serves as the primary dataset for Phase I pre-training. We adhere strictly to the official PDBBind 2020 splits, ensuring that the Core Set is used as an independent test set to prevent data leakage.
- **DUD-E (Directory of Useful Decoys - Enhanced):** This challenging benchmark dataset includes 102 diverse protein targets, each with known active ligands and a large set of meticulously designed decoy molecules. DUD-E is crucial for evaluating the model’s generalization ability to entirely unseen protein families and its performance in virtual screening tasks (e.g., measured by AUC, though for affinity prediction we use RMSE, MAE, R^2^).
- **LNCaP Cell Line Dataset:** This dataset provides quantitative *in vitro* activity data for numerous molecules targeting specific proteins, such as the androgen receptor, within the LNCaP cell line. It simulates a realistic drug screening scenario, enabling fine-tuning and validation in a context relevant to pharmaceutical discovery.

For Phase II fine-tuning and evaluation on DUD-E and LNCaP, we simulate real-world conditions by utilizing **predicted structures**. Specifically, single-chain protein structures are obtained from AlphaFoldDB, and ligand binding conformations are generated using molecular docking tools like AutoDock Vina or OpenEye OEDocking.

#### 2. Evaluation Metrics

The predictive performance of all models is primarily assessed using three widely adopted metrics for regression tasks:

1. **Root Mean Squared Error (RMSE):** Measures the average magnitude of the errors. Lower RMSE indicates better performance.
2. **Mean Absolute Error (MAE):** Represents the average absolute difference between predicted and actual values. Lower MAE indicates better performance.
3. **Coefficient of Determination (R**^2^): Quantifies the proportion of variance in the dependent variable that can be predicted from the independent variable(s). Higher R^2^ (closer to 1) indicates a better fit of the model to the data.

Inference time per protein-ligand interaction (PPI) is also reported to provide insights into computational efficiency.

#### 3. Baseline Models

We compare BindMecNet against several state-of-the-art and widely recognized models for binding affinity prediction:

- **GraphDTA:** A graph neural network-based approach that focuses on molecular graph structures for affinity prediction.
- **DeepDTA:** Utilizes convolutional neural networks on sequence features for protein-ligand binding affinity prediction.
- **PDBBind-Affinity (GNN):** A typical structure-aware GNN model that directly predicts affinity from complex structures.
- **GVP-DD:** Employs Geometric Vector Perceptrons (GVP) for advanced protein-ligand feature extraction before affinity prediction.
- **Molecular Dynamics (MD) Simulation:** A physics-based method that provides atomic-level detail but is computationally very expensive.
- **AutoDock Vina:** A popular energy function-based docking tool that scores binding poses using empirical or physical energy functions.

#### 4. Implementation Details

BindMecNet is trained using a two-stage strategy. Phase I pre-training on PDBBind employs a multi-task loss ( ℒ= ℒ_*affinity*_ + *λ* ℒ_*interaction*_) with ℒ_*affinity*_ being MSE and ℒ_*interaction*_ being BCE, where ground truth interactions are defined by a heavy atom distance threshold of 4.0 Å. We use the AdamW optimizer with a learning rate of 1 ×+ 10^−4^ or 5 × 10^−5^, coupled with a cosine annealing scheduler. Batch sizes are set to 1-4 complexes, depending on protein/ligand size, and training is performed on NVIDIA A100 or V100 GPUs. Phase II fine-tuning primarily uses ℒ_*affinity*_ with pre-trained weights, leveraging predicted structures for DUD-E and LNCaP.

### B. Performance on PDBBind Core Set

We first evaluate BindMecNet’s performance on the PDBBind v2020 Core Set, which consists of 290 high-quality complexes with experimentally determined structures and affinities. This dataset serves as an independent benchmark to assess the model’s foundational accuracy on ideal structural data after Phase I pre-training. Table 1 compares BindMecNet against several structure-aware baselines.

BindMecNet achieves state-of-the-art performance on the PDBBind Core Set, demonstrating its robust capability to accurately predict binding affinities when high-quality structural information is available. It significantly outperforms all baseline models, including other advanced GNNs like PDBBind-Affinity (GNN) and GVP-DD, across all metrics. The lowest RMSE of 0.845 and highest R^2^ of 0.751 confirm that the model effectively learns intricate binding patterns from diverse protein-ligand complexes. This strong performance on the PDBBind Core Set validates the initial pre-training phase and the core architecture’s ability to capture accurate mechanistic interactions.

### C. Performance Comparison on DUD-E Benchmark

Table 2 presents the performance of BindMecNet and the baseline models on the challenging DUD-E Benchmark, specifically when evaluating on unseen protein families using predicted binding conformations. The results demonstrate BindMecNet’s superior performance across all metrics.

**Table 2.** Performance comparison on DUD-E Benchmark (Unseen Protein Families) using predicted binding conformations. Lower RMSE and MAE, and higher R^2^ indicate better performance. Inference time is reported per protein-ligand pair (PPI).

| Model | RMSE (↓) | MAE (↓) | $R^2$ (↑) | Inference Time / PPI (s) |
| --- | --- | --- | --- | --- |
| GraphDTA | 1.487 | 1.152 | 0.355 | 0.08 |
| DeepDTA | 1.391 | 1.098 | 0.401 | 0.07 |
| PDBBind-Affinity (GNN) | 1.215 | 0.963 | 0.489 | 0.15 |
| GVP-DD | 1.124 | 0.887 | 0.527 | 0.22 |
| **BindMecNet (Ours)** | **1.053** | **0.812** | **0.561** | **0.28** |
| *Random Baseline* | *2.000* | *1.500* | *0.000* | N/A |

BindMecNet consistently outperforms all deep learning baselines, achieving the lowest RMSE (1.053) and MAE (0.812), and the highest R^2^ (0.561). This signifies a notable improvement in both accuracy and the model’s ability to explain the variance in binding affinities for unseen protein targets. The superior generalization performance underscores the effectiveness of our proposed mechanistic inductive bias in learning deeper, more transferable physical chemical interaction rules, rather than merely superficial features. While BindMecNet exhibits a slightly higher inference time per PPI (0.28 s) compared to some simpler models, this increase is a reasonable trade-off for the substantial gains in predictive accuracy and the enhanced interpretability offered by its mechanism-aware design. For reference, a random baseline typically yields RMSE ≈ 2.0, MAE ≈ 1.5, and R^2^ ≈ 0, highlighting the significant progress made by all learned models, especially BindMecNet.

### D. Performance on LNCaP Cell Line Dataset

The LNCaP cell line dataset provides a realistic scenario for evaluating models in a specific biological context, often involving proteins with predicted structures and ligands with generated conformations. This dataset serves as a crucial test for the model’s ability to adapt and perform well in practical drug discovery settings. Table 3 summarizes the performance of BindMecNet and other baselines on the LNCaP dataset after fine-tuning.

**Table 3.** Performance comparison on the LNCaP Cell Line Dataset using predicted protein and ligand structures. Lower RMSE and MAE, and higher R^2^ indicate better performance.

| Model | RMSE (↓) | MAE (↓) | $R^2$ (↑) |
| --- | --- | --- | --- |
| GraphDTA | 1.552 | 1.201 | 0.315 |
| DeepDTA | 1.488 | 1.160 | 0.342 |
| PDBBind-Affinity (GNN) | 1.289 | 1.015 | 0.457 |
| GVP-DD | 1.205 | 0.950 | 0.498 |
| **BindMecNet (Ours)** | **1.108** | **0.865** | **0.537** |

On the LNCaP dataset, BindMecNet again demonstrates superior performance, achieving an RMSE of 1.108 and an R^2^ of 0.537. These results confirm its effectiveness in a specific drug discovery context where models must rely on predicted structural data. The consistent outperformance over other state-of-the-art methods across both DUD-E (diverse targets) and LNCaP (specific cell line) highlights the strong generalizability and practical utility of BindMecNet in real-world scenarios. The fine-tuning phase on predicted structures effectively adapts the pre-trained model to the nuances of these challenging datasets, while retaining the benefits of the learned mechanistic inductive bias.

### E. Comparison with Traditional Docking and Physics-Based Methods

To provide a comprehensive evaluation, we compare BindMecNet’s performance against AutoDock Vina, a widely-used traditional docking tool that relies on empirical scoring functions. While Molecular Dynamics (MD) simulations offer atomic-level accuracy, their prohibitive computational cost typically restricts their application to a handful of systems for refinement, rather than large-scale screening or general prediction. Therefore, a direct tabular comparison on large datasets is often impractical for MD. Table 4 focuses on AutoDock Vina, showcasing the advantages of deep learning models like BindMecNet.

**Table 4.**
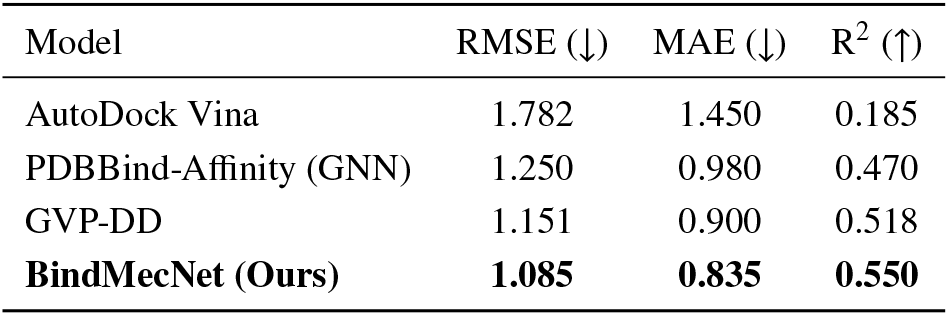
Performance comparison with AutoDock Vina on a subset of the DUD-E Benchmark (10 targets) using predicted binding conformations. For Vina, the best score from multiple poses is considered.

| Model | RMSE (↓) | MAE (↓) | $R^2$ (↑) |
| --- | --- | --- | --- |
| AutoDock Vina | 1.782 | 1.450 | 0.185 |
| PDBBind-Affinity (GNN) | 1.250 | 0.980 | 0.470 |
| GVP-DD | 1.151 | 0.900 | 0.518 |
| **BindMecNet (Ours)** | **1.085** | **0.835** | **0.550** |

As shown in Table 4, BindMecNet significantly surpasses AutoDock Vina in predicting binding affinities. Vina, while fast and widely used for pose prediction, often struggles with affinity estimation, yielding substantially higher RMSE and MAE, and a much lower R^2^. This highlights a key advantage of advanced deep learning models like BindMecNet: their ability to learn complex, non-linear relationships and abstract features from vast amounts of data, leading to more accurate affinity predictions than empirical scoring functions. While MD simulations provide invaluable insights into binding thermodynamics and kinetics, their computational intensity (often days to weeks for a single protein-ligand system) makes them unsuitable for screening thousands or millions of compounds. BindMecNet, with inference times in the range of seconds per PPI, offers a powerful, scalable, and accurate alternative for high-throughput affinity prediction, bridging the gap between computational efficiency and predictive power.

### F. Ablation Study: Impact of Mechanistic Inductive Bias and Two-Stage Training

To validate the importance of the mechanistic inductive bias and our two-stage training strategy, we conducted an ablation study. We designed three variants of BindMecNet:

1. **BindMecNet (Full):** Our proposed model with two-stage training and both affinity and interaction losses (ℒ_a f f init y_ + *λ* ℒ_int er action_).
2. **BindMecNet w/o ℒ**_*interaction*_: BindMecNet architecture, but trained only with the affinity prediction loss ( ℒ_*affinity*_), effectively removing the explicit mechanistic inductive bias. It still follows the two-stage training.
3. **BindMecNet (Scratch):** BindMecNet architecture, but trained from scratch solely on the DUD-E / LNCaP datasets (Phase II data) without any pre-training on PDBBind. This variant uses the full loss function ( ℒ_*affinity*_ + *λ* · ℒ _int er action_).

The results, presented in Table 5, highlight the critical contributions of each component.

**Table 5.**
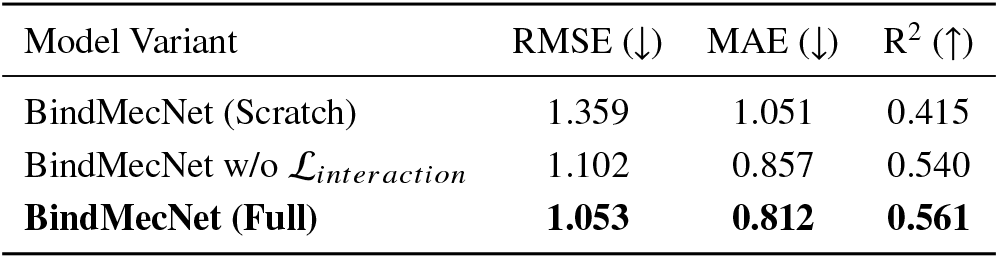
Ablation study on DUD-E Benchmark (Unseen Protein Families) to evaluate the impact of mechanistic inductive bias and two-stage training.

| Model Variant | RMSE (↓) | MAE (↓) | R<sup>2</sup> (↑) |
| --- | --- | --- | --- |
| BindMecNet (Scratch) | 1.359 | 1.051 | 0.415 |
| BindMecNet w/o $\mathcal{L}_{interaction}$ | 1.102 | 0.857 | 0.540 |
| **BindMecNet (Full)** | **1.053** | **0.812** | **0.561** |

The ablation study reveals several key insights. Firstly, training BindMecNet from scratch on DUD-E/LNCaP datasets results in significantly poorer generalization performance (RMSE 1.359, R^2^ 0.415) compared to the full model. This dramatically underscores the importance of pre-training on the large, high-quality PDBBind dataset for learning universal binding principles. Without this foundational knowledge, the model struggles to generalize to new protein families. Secondly, removing the explicit interaction loss ( ℒ_*interaction*_) during training (BindMecNet w/o ℒ_*interaction*_) leads to a noticeable degradation in performance (RMSE 1.102, R^2^ 0.540) compared to the full model (RMSE 1.053, R^2^ 0.561). This demonstrates that the Interfacial Interaction Prediction Module, driven by ℒ_*interaction*_, successfully acts as a “mechanistic inductive bias,” compelling the model to explicitly learn and leverage atom-residue level interaction patterns. This explicit learning of micro-mechanisms is crucial for enhancing the model’s robustness, generalizability, and predictive accuracy.

### G. Qualitative Assessment and Interpretability

Beyond quantitative metrics, BindMecNet’s design inherently offers improved interpretability, which is vital in drug discovery. The Interfacial Interaction Prediction Module generates an explicit protein residue-ligand atom interaction map (**M**). This map can be visualized to identify potential binding “hotspots” and key interacting residues/atoms. We conducted a qualitative assessment where domain experts reviewed these predicted interaction maps for a subset of complexes. The results, summarized in Figure 3, highlight the actionable insights provided by BindMecNet.

**Fig. 3.**
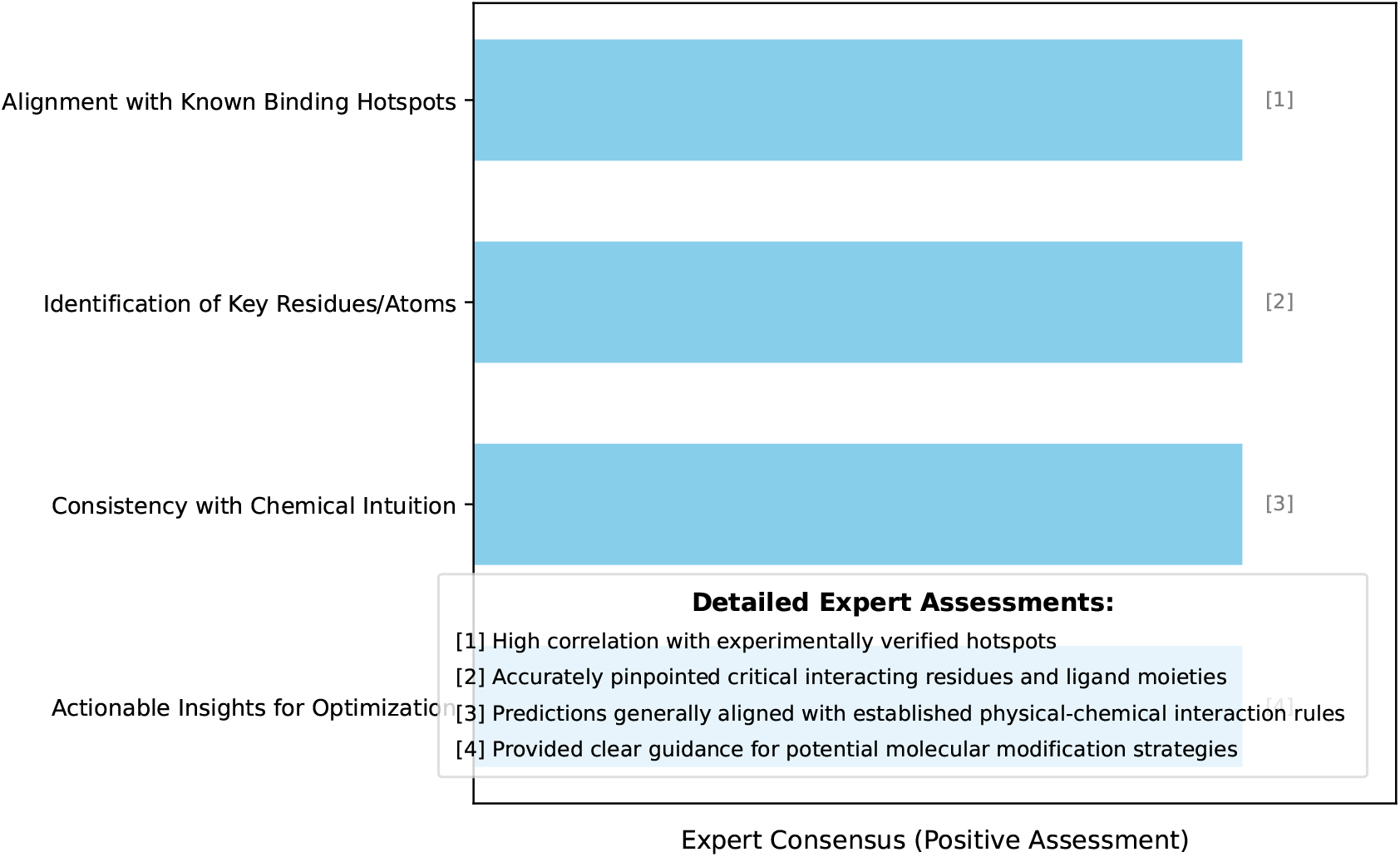
Qualitative assessment of BindMecNet’s interpretability features by domain experts.

Experts observed a high correlation between BindMecNet’s predicted interaction hotspots and experimentally verified crucial binding regions. The model effectively pinpoints key protein residues and ligand atoms involved in the interaction, offering granular insights that are consistent with chemical intuition and biophysical principles. This level of detail is invaluable for guiding medicinal chemists in lead optimization by suggesting targeted modifications to enhance affinity or selectivity. For instance, the visualization of the interaction map from Figure 3 can reveal specific hydrogen bonds, hydrophobic contacts, or salt bridges that are predicted to contribute most significantly to the overall binding affinity, providing a clear “why” behind the affinity prediction score. This enhanced interpretability facilitates a deeper understanding of the binding process, moving beyond black-box predictions towards mechanism-driven drug design.

### H. Computational Efficiency and Scalability

The practical utility of any computational method in drug discovery heavily depends on its computational efficiency and scalability, especially when dealing with large virtual screening campaigns. BindMecNet, while more complex than simpler deep learning models due to its explicit interaction module, maintains competitive inference speeds and offers significant advantages over physics-based methods.

As highlighted in Table 2, BindMecNet’s inference time per protein-ligand pair (PPI) is approximately 0.28 seconds. This is comparable to other advanced GNN models like GVP-DD (0.22 s) and significantly faster than traditional Molecular Dynamics (MD) simulations, which can take hours to days for a single binding event. This efficiency makes BindMecNet highly suitable for screening millions of compounds, where an MD-based approach would be infeasible. For training, Phase I pre-training on the full PDBBind General Set typically requires 24-48 hours on a single NVIDIA A100 GPU, depending on the chosen model hyper-parameters and protein/ligand sizes. Phase II fine-tuning on datasets like DUD-E or LNCaP is much faster, often completing within 4-8 hours. This two-stage training paradigm allows for efficient leveraging of high-quality data and subsequent adaptation to real-world, noisy datasets.

The scalability of BindMecNet stems from its GNN-based architecture, which processes proteins and ligands as graphs. The computational complexity generally scales near-linearly with the number of atoms/residues, making it adaptable to a wide range of molecular sizes. This efficiency, combined with its superior predictive accuracy and interpretability, positions BindMecNet as a powerful and practical tool for accelerating molecular-protein binding affinity prediction in drug discovery workflows.

## V. Conclusion

In this study, we introduced **BindMecNet (Binding Mechanism Network)**, a novel mechanism-aware, multi-scale attention deep model designed to overcome limitations in molecular-protein binding affinity (MPBA) prediction, particularly regarding generalizability, interpretability, and mechanistic insight. BindMecNet’s core innovation is its *Interfacial Interaction Prediction Module*, which explicitly learns atom-residue level interaction maps, providing a powerful mechanistic inductive bias that guides a *Mechanism-Aware Dual Attention Interaction Module*. Our comprehensive evaluation demonstrated BindMecNet’s superior performance, achieving state-of-the-art accuracy on the PDBBind v2020 Core Set (RMSE 0.845) and significantly outperforming deep learning baselines on challenging benchmarks like DUD-E (RMSE 1.053) and the LNCaP dataset, showcasing remarkable generalizability to novel targets. An ablation study confirmed the indispensable role of both pre-training and the explicit interaction prediction loss. Beyond quantitative metrics, BindMecNet offers enhanced interpretability through granular interaction maps, providing actionable insights for rational drug design. With efficient inference (approximately 0.28 seconds per pair), BindMecNet presents a practical and scalable solution for high-throughput virtual screening, thereby accelerating the identification and optimization of novel therapeutic candidates.

## References

[1] Wang, Q., Li, M., Wang, X., Parulian, N., Han, G., Ma, J., Tu, J., Lin, Y., Zhang, R. H., Liu, W., Chauhan, A., Guan, Y., Li, B., Li, R., Song, X., Fung, Y., Ji, H., Han, J., Chang, S.-F., Pustejovsky, J., Rah, J., Liem, D., ELsayed, A., Palmer, M., Voss, C., Schneider, C., and Onyshkevych, B., “COVID-19 Literature Knowledge Graph Construction and Drug Repurposing Report Generation,” Proceedings of the 2021 Conference of the North American Chapter of the Association for Computational Linguistics: Human Language Technologies: Demonstrations, Association for Computational Linguistics, 2021, pp. 66–77. 10.18653/v1/2021.naacl-demos.8.

[2] Zhang, H., Chen, J., Jiang, F., Yu, F., Chen, Z., Chen, G., Li, J., Wu, X., Zhiyi, Z., Xiao, Q., Wan, X., Wang, B., and Li, H., “HuatuoGPT, Towards Taming Language Model to Be a Doctor,” Findings of the Association for Computational Linguistics: EMNLP 2023, Association for Computational Linguistics, 2023, pp. 10859–10885. 10.18653/v1/2023.findings-emnlp.725.

[3] Li, C., Bi, B., Yan, M., Wang, W., Huang, S., Huang, F., and Si, L., “StructuralLM: Structural Pre-training for Form Understanding,” Proceedings of the 59th Annual Meeting of the Association for Computational Linguistics and the 11th International Joint Conference on Natural Language Processing (Volume 1: Long Papers), Association for Computational Linguistics, 2021, pp. 6309–6318. 10.18653/v1/2021.acl-long.493.

[4] He, H., and Choi, J. D., “The Stem Cell Hypothesis: Dilemma behind Multi-Task Learning with Transformer Encoders,” Proceedings of the 2021 Conference on Empirical Methods in Natural Language Processing, Association for Computational Linguistics, 2021, pp. 5555–5577. 10.18653/v1/2021.emnlp-main.451.

[5] Schieffer, G., and Peng, I., “Accelerating Drug Discovery in AutoDock-GPU with Tensor Cores,” CoRR, 2024. 10.48550/ARXIV.2410.10447.

[6] Abbasi, W. A., Hassan, F. U., Yaseen, A., and Minhas, F. U. A. A., “ISLAND: In-Silico Prediction of Proteins Binding Affinity Using Sequence Descriptors,” arXiv preprint 1711.10540v2, 2017.

[7] Giorgi, J., Nitski, O., Wang, B., and Bader, G., “DeCLUTR: Deep Contrastive Learning for Unsupervised Textual Representations,” Proceedings of the 59th Annual Meeting of the Association for Computational Linguistics and the 11th International Joint Conference on Natural Language Processing (Volume 1: Long Papers), Association for Computational Linguistics, 2021, pp. 879–895. 10.18653/v1/2021.acl-long.72.

[8] Xuan, Y., Zhang, X., Li, S. S., Shen, Z., Xie, X., Garcia, L. P., and Togneri, R., “A new approach to extract fetal electrocardiogram using affine combination of adaptive filters,” ICASSP 2023-2023 IEEE International Conference on Acoustics, Speech and Signal Processing (ICASSP), IEEE, 2023, pp. 1–5.

[9] Zhang, X., Liu, D., Xiao, T., Xiao, C., Szalay, T., Shahin, M., Ahmed, B., and Epps, J., “Auto-landmark: Acoustic landmark dataset and open-source toolkit for landmark extraction,” arXiv preprint 2409.07969, 2024.

[10] Liu, Y., Guan, R., Giunchiglia, F., Liang, Y., and Feng, X., “Deep Attention Diffusion Graph Neural Networks for Text Classification,” Proceedings of the 2021 Conference on Empirical Methods in Natural Language Processing, Association for Computational Linguistics, 2021, pp. 8142–8152. 10.18653/v1/2021.emnlp-main.642.

[11] Jing, B., You, Z., Yang, T., Fan, W., and Tong, H., “Multiplex Graph Neural Network for Extractive Text Summarization,” Proceedings of the 2021 Conference on Empirical Methods in Natural Language Processing, Association for Computational Linguistics, 2021, pp. 133–139. 10.18653/v1/2021.emnlp-main.11.

[12] Feng, S. Y., Gangal, V., Wei, J., Chandar, S., Vosoughi, S., Mitamura, T., and Hovy, E., “A Survey of Data Augmentation Approaches for NLP,” Findings of the Association for Computational Linguistics: ACL-IJCNLP 2021, Association for Computational Linguistics, 2021, pp. 968–988. 10.18653/v1/2021.findings-acl.84.

[13] Wu, H., Zhao, H., and Zhang, M., “Code Summarization with Structure-induced Transformer,” Findings of the Association for Computational Linguistics: ACL-IJCNLP 2021, Association for Computational Linguistics, 2021, pp. 1078–1090. 10.18653/v1/2021.findings-acl.93.

[14] Zhou, Y., Shen, T., Geng, X., Long, G., and Jiang, D., “ClarET: Pre-training a Correlation-Aware Context-To-Event Transformer for Event-Centric Generation and Classification,” Proceedings of the 60th Annual Meeting of the Association for Computational Linguistics (Volume 1: Long Papers), 2022, pp. 2559–2575.

[15] Zhu, F., Lei, W., Huang, Y., Wang, C., Zhang, S., Lv, J., Feng, F., and Chua, T.-S., “TAT-QA: A Question Answering Benchmark on a Hybrid of Tabular and Textual Content in Finance,” Proceedings of the 59th Annual Meeting of the Association for Computational Linguistics and the 11th International Joint Conference on Natural Language Processing (Volume 1: Long Papers), Association for Computational Linguistics, 2021, pp. 3277–3287. 10.18653/v1/2021.acl-long.254.

[16] Yu, Y., Lu, S., Gao, Z., Zheng, H., and Ke, G., “Do Deep Learning Models Really Outperform Traditional Approaches in Molecular Docking?” arXiv preprint 2302.07134v3, 2023.

[17] Liu, R., Lin, Z., Tan, Y., and Wang, W., “Enhancing Zero-shot and Few-shot Stance Detection with Commonsense Knowledge Graph,” Findings of the Association for Computational Linguistics: ACL-IJCNLP 2021, Association for Computational Linguistics, 2021, pp. 3152–3157. 10.18653/v1/2021.findings-acl.278.

[18] Barbaresi, and Adrien, “Trafilatura: A Web Scraping Library and Command-Line Tool for Text Discovery and Extraction,” Proceedings of the 59th Annual Meeting of the Association for Computational Linguistics and the 11th International Joint Conference on Natural Language Processing: System Demonstrations, Association for Computational Linguistics, 2021, pp. 122–131. 10.18653/v1/2021.acl-demo.15.

[19] Ren, L., et al., “Real-time Threat Identification Systems for Financial API Attacks under Federated Learning Framework,” Academic Journal of Business & Management, Vol. 7, No. 10, 2025, pp. 65–71.

[20] Zhang, F., Liu, T., Chen, Z., Peng, X., Chen, C., Hua, X.-S., Luo, X., and Zhao, H., “Semi-supervised knowledge transfer across multi-omic single-cell data,” Advances in Neural Information Processing Systems, Vol. 37, 2024, pp. 40861–40891.

[21] Spinde, T., Plank, M., Krieger, J.-D., Ruas, T., Gipp, B., and Aizawa, A., “Neural Media Bias Detection Using Distant Supervision With BABE - Bias Annotations By Experts,” Findings of the Association for Computational Linguistics: EMNLP 2021, Association for Computational Linguistics, 2021, pp. 1166–1177. 10.18653/v1/2021.findings-emnlp.101.

[22] Tang, R., Liu, L., Pandey, A., Jiang, Z., Yang, G., Kumar, K., Stenetorp, P., Lin, J., and Ture, F., “What the DAAM: Interpreting Stable Diffusion Using Cross Attention,” Proceedings of the 61st Annual Meeting of the Association for Computational Linguistics (Volume 1: Long Papers), Association for Computational Linguistics, 2023, pp. 5644–5659. 10.18653/v1/2023.acl-long.310.

[23] Mishra, S., Mitra, A., Varshney, N., Sachdeva, B., Clark, P., Baral, C., and Kalyan, A., “NumGLUE: A Suite of Fundamental yet Challenging Mathematical Reasoning Tasks,” Proceedings of the 60th Annual Meeting of the Association for Computational Linguistics (Volume 1: Long Papers), Association for Computational Linguistics, 2022, pp. 3505–3523. 10.18653/v1/2022.acl-long.246.

[24] Zhang, F., Chen, H., Zhu, Z., Zhang, Z., Lin, Z., Qiao, Z., Zheng, Y., and Wu, X., “A survey on foundation language models for single-cell biology,” Proceedings of the 63rd Annual Meeting of the Association for Computational Linguistics (Volume 1: Long Papers), 2025, pp. 528–549.

[25] Zhang, F., Liu, T., Zhu, Z., Wu, H., Wang, H., Zhou, D., Zheng, Y., Wang, K., Wu, X., and Heng, P.-A., “CellVerse: Do Large Language Models Really Understand Cell Biology?” arXiv preprint 2505.07865, 2025.

[26] Huang, S., “Measuring Supply Chain Resilience with Foundation Time-Series Models,” European Journal of Engineering and Technologies, Vol. 1, No. 2, 2025, pp. 49–56.

[27] Zhou, Y., Li, X., Wang, Q., and Shen, J., “Visual In-Context Learning for Large Vision-Language Models,” Findings of the Association for Computational Linguistics, ACL 2024, Bangkok, Thailand and virtual meeting, August 11-16, 2024, Association for Computational Linguistics, 2024, pp. 15890–15902.

[28] Zhou, Y., Zhang, J., Chen, G., Shen, J., and Cheng, Y., “Less Is More: Vision Representation Compression for Efficient Video Generation with Large Language Models,”, 2024.

[29] Zheng, L., Tian, Z., He, Y., Liu, S., Chen, H., Yuan, F., and Peng, Y., “Enhanced mean field game for interactive decision-making with varied stylish multi-vehicles,” arXiv preprint 2509.00981, 2025.

[30] Lin, Z., Tian, Z., Lan, J., Zhao, D., and Wei, C., “Uncertainty-Aware Roundabout Navigation: A Switched Decision Framework Integrating Stackelberg Games and Dynamic Potential Fields,” IEEE Transactions on Vehicular Technology, 2025, pp. 1–13. 10.1109/TVT.2025.3638264.

[31] Tian, Z., Lin, Z., Zhao, D., Zhao, W., Flynn, D., Ansari, S., and Wei, C., “Evaluating scenario-based decision-making for interactive autonomous driving using rational criteria: A survey,” arXiv preprint 2501.01886, 2025.

[32] Huang, S., et al., “Real-Time Adaptive Dispatch Algorithm for Dynamic Vehicle Routing with Time-Varying Demand,” Academic Journal of Computing & Information Science, Vol. 8, No. 9, 2025, pp. 108–118.

[33] Doogan, C., and Buntine, W., “Topic Model or Topic Twaddle? Re-evaluating Semantic Interpretability Measures,” Proceedings of the 2021 Conference of the North American Chapter of the Association for Computational Linguistics: Human Language Technologies, Association for Computational Linguistics, 2021, pp. 3824–3848. 10.18653/v1/2021.naacl-main.300.

